## Supplementary Information for "Coevolution of male and female mate choice can destabilize reproductive isolation"

##### Contents

Pages 2 - 15: Supplementary Figs. 1-15

Pages 16 - 17: Supplementary Note 1: Cost of female choosiness

Pages 18 - 19: Supplementary Note 2: Continuous choosiness

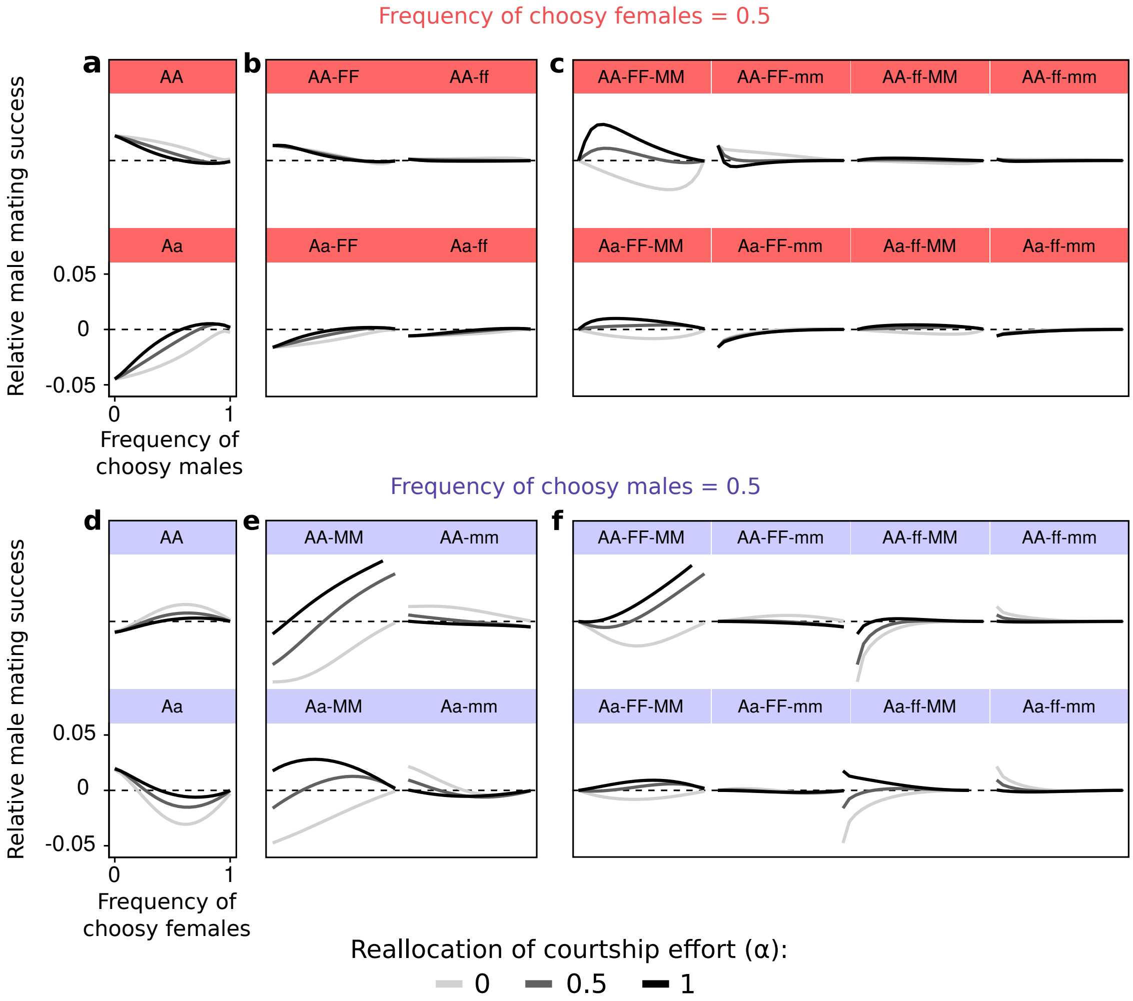


**Supplementary Figure 1:** Relative male mating success (compared to the mating success expected under random mating) for $s = 0.2$ with varying frequencies of choosy males (a, b, c) or choosy females (d, e, f) and different $\alpha$ values. Relative male mating successes involving genotype aa are similar to the ones involving genotypes AA and are not represented. Likewise, relative male mating successes involving heterozygous choosiness genotypes Ff or Mm reflect the ones involving homozygous choosiness genotypes ff and mm, and are not shown either. The scale is the same for all subfigures. Relative male mating success of genotype $x$ is calculated as $\sum_f [M_{x,f} - p_f.p_x]$ with the frequencies at ecological equilibrium. Sexual selection acts through variations in male mating success among genotypes (females' mating success is equal in our model). Here, if males are not choosy, homozygous males with genotype AA or aa have a high mating success (a); female choosiness generates positive frequency-dependent sexual selection that favours the most common ecological genotypes (sexual selection represented by the green arrow in Fig. 1). On the contrary, if females are not choosy and $\alpha = 1$, homozygous males with genotype AA or aa have a low mating success (d). Because choosy males are mostly homozygous at the $A$ locus (linkage disequilibrium; Supplementary Fig. 2), competition for access to homozygous females is stiffer than for access to heterozygous females. Choosy homozygous males (AA-MM and aa-MM) court almost exclusively matching females and therefore place themselves in a disadvantageous competitive setting lowering their mating success (e). Therefore, sexual selection caused by distorted male-male competition directly inhibits the evolution of male choosiness (pink arrow in Fig. 1). If $\alpha < 1$, lost courtship opportunities caused by male choosiness also decrease the mating success of choosy males (e) (blue arrow in Fig. 1). This selection pressure is independent from the one caused by distorted male-male competition; all choosy males (not only AA-MM and aa-MM males) have a lower mating success if $\alpha$ is lower (e). Finally, for all $\alpha$, female choosiness increases the mating success of choosy males because choosy females reject nonmatching males that are mainly nonchoosy (e) (red arrow in Fig. 1). This selection pressure is independent from indirect sexual selection induced by female choosiness (green arrow in Fig. 1); all choosy males (not only AA-MM and aa-MM males) have a higher mating success if the frequency of choosy females is higher (e).

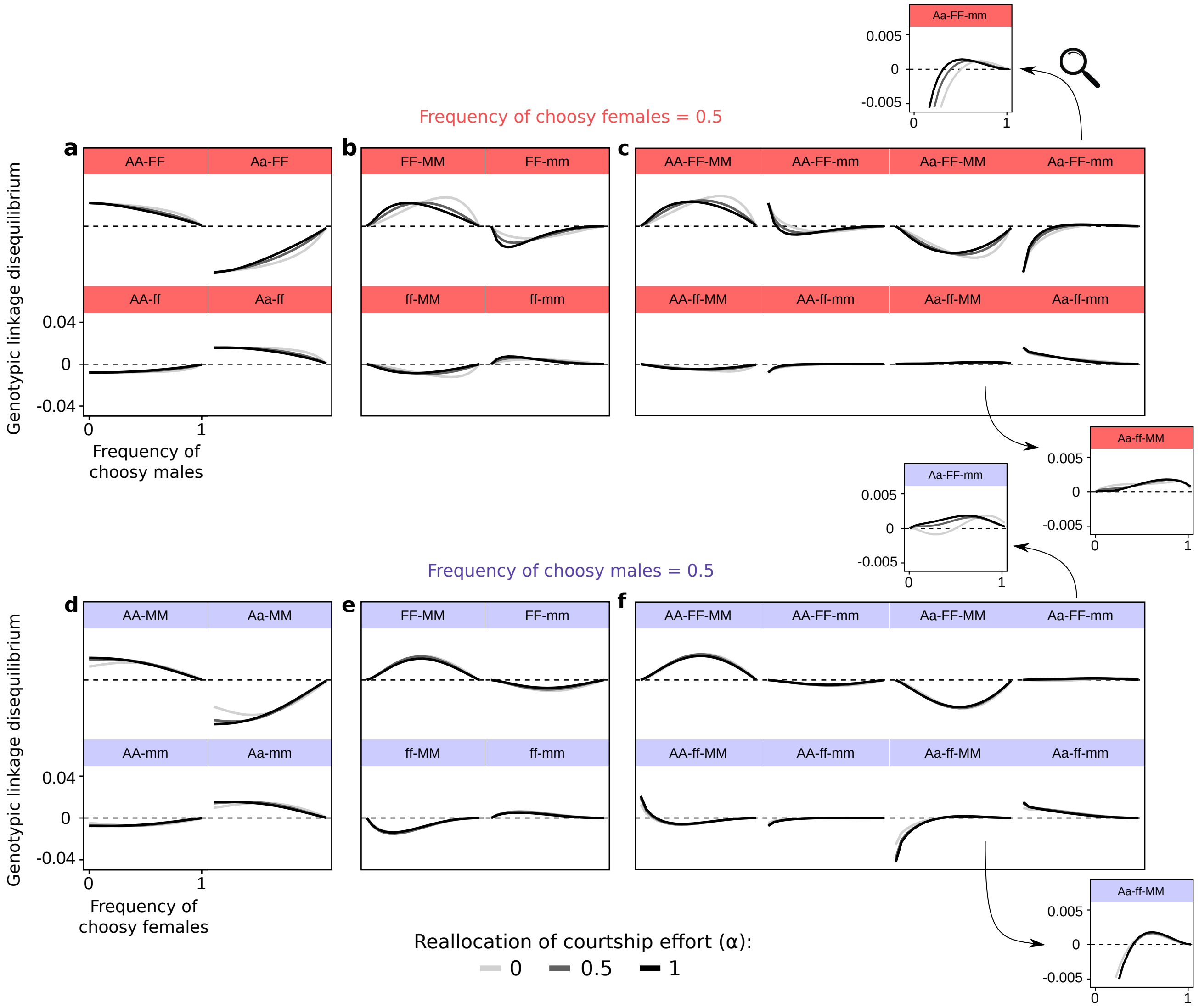


**Supplementary Figure 2:** Genotypic linkage disequilibrium for $s = 0.2$ with varying frequencies of choosy males (a, b, c) or choosy females (d, e, f) and different $\alpha$ values. Linkage disequilibria involving genotype aa are similar to the ones involving genotype AA and are not represented. Likewise, linkage disequilibria involving heterozygous choosiness genotypes Ff or Mm reflect the ones involving homozygous choosiness genotypes ff and mm, and are not shown either. The scale is the same for all subfigures. We calculate the linkage disequilibrium between two diploid genotypes $x$ and $y$ at different loci (e.g., between genotypes FF and AA) as: $D_{x-y} = \text{freq}_{xy} - \text{freq}_x.\text{freq}_y$. $D_{x-y} > 0$ indicates a positive linkage disequilibrium between genotypes $x$ and $y$. Likewise, we calculate the linkage disequilibrium between three diploid genotypes $x$, $y$ and $z$ at different loci as: $D_{x-y-z} = \text{freq}_{xyz} - \text{freq}_x.\text{freq}_y.\text{freq}_z$. Selection causes linkage disequilibrium among loci. This linkage disequilibrium is qualitatively similar for all $\alpha$ and $s$ values tested (not shown). Here, female and male choosiness genotypes are linked to homozygous genotypes at the $A$ locus ($D_{AA-FF}$, $D_{aa-FF}$, $D_{AA-MM}$ and $D_{aa-MM} > 0$, in subfigures a and d). Indirect viability and sexual selection acting on the $A$ locus can therefore indirectly favour the evolution of female and male choosiness (indirect selection represented by the dashed black and green arrows in Fig. 1). However these positive linkage disequilibria vanish if the other sex is choosy; indirect selection is weak under these conditions. Additionally, female and male choosiness genotypes are associated ($D_{FF-MM} > 0$ in subfigures b and e), especially in homozygotes at the ecological locus ($D_{AA-FF-MM} > 0$, $D_{aa-FF-MM} > 0$ and $D_{Aa-FF-MM} < 0$ in subfigures c and f). On the contrary, genotypes ff-MM and FF-mm are somewhat more likely to be heterozygotes Aa ($D_{Aa-ff-MM} > 0$, $D_{Aa-FF-mm} > 0$ in subfigures c and f, c.f. zoom of the subplots). While choosy heterozygous males benefit from reduced male-male competition (increasing the frequency of alleles coding for male choosiness; light blue in Fig. 2), choosy homozygous males (which are strongly linked to female choosiness) face intensified male-male competition (reducing the frequency of alleles coding for female choosiness; dark red in Fig. 2). This is why sexual selection acting on the $M$ locus can indirectly affect the evolution of female choosiness (indirect selection represented by the dashed pink, red and blue arrows in Fig. 1).

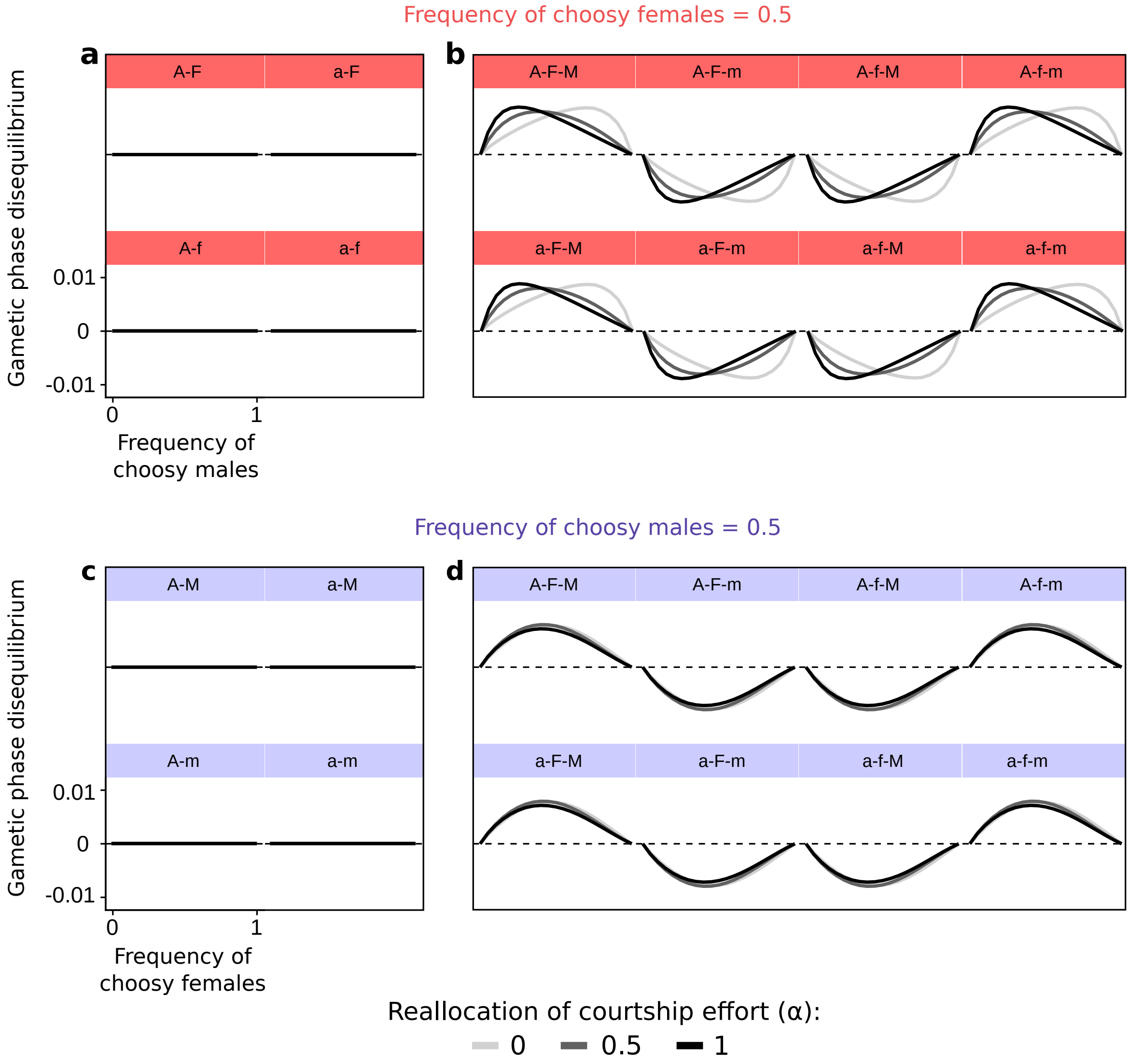


**Supplementary Figure 3:** Gametic phase disequilibrium for $s = 0.2$ with varying frequencies of choosy males (a, b) or choosy females (c, d) and different $\alpha$ values. The scale is the same for all subfigures. We calculate the gametic phase disequilibrium between alleles $x$ and $y$ at different loci (e.g., between alleles F and A) as: $D'_{x-y} = \text{freq}'_{xy} - \text{freq}'_x.\text{freq}'_y$, with freq$'$ referring to frequencies in haploid gametes. $D'_{x-y} > 0$ indicates a positive linkage disequilibrium between alleles $x$ and $y$ in haploid gametes. Likewise, we calculate the gametic phase disequilibrium between three alleles $x$, $y$ and $z$ at different loci as: $D'_{x-y-z} = \text{freq}'_{xyz} - \text{freq}'_x.\text{freq}'_y.\text{freq}'_z$. Selection causes gametic phase disequilibrium among loci. This gametic phase disequilibrium is qualitatively similar for all $\alpha$ and $s$ values tested (not shown). In gametes, alleles coding for choosiness (M and F) are not linked to a particular allele (A or a) at the ecological locus ($D'_{A-F} = D'_{a-F} = D'_{A-M} = D'_{a-M} = 0$ in subfigures a and c) because they associate with both homozygous ecotypes (genotypes AA and aa; Supplementary Fig. 2). Nonetheless, alleles coding for choosiness associate with each other in gametes ($D'_{A-F-M} > 0$ and $D'_{a-F-M} > 0$ in subfigures b and d). This occurs because genotypes FF and MM associate within both ecotypes (genotypes AA and aa; Supplementary Fig. 2).

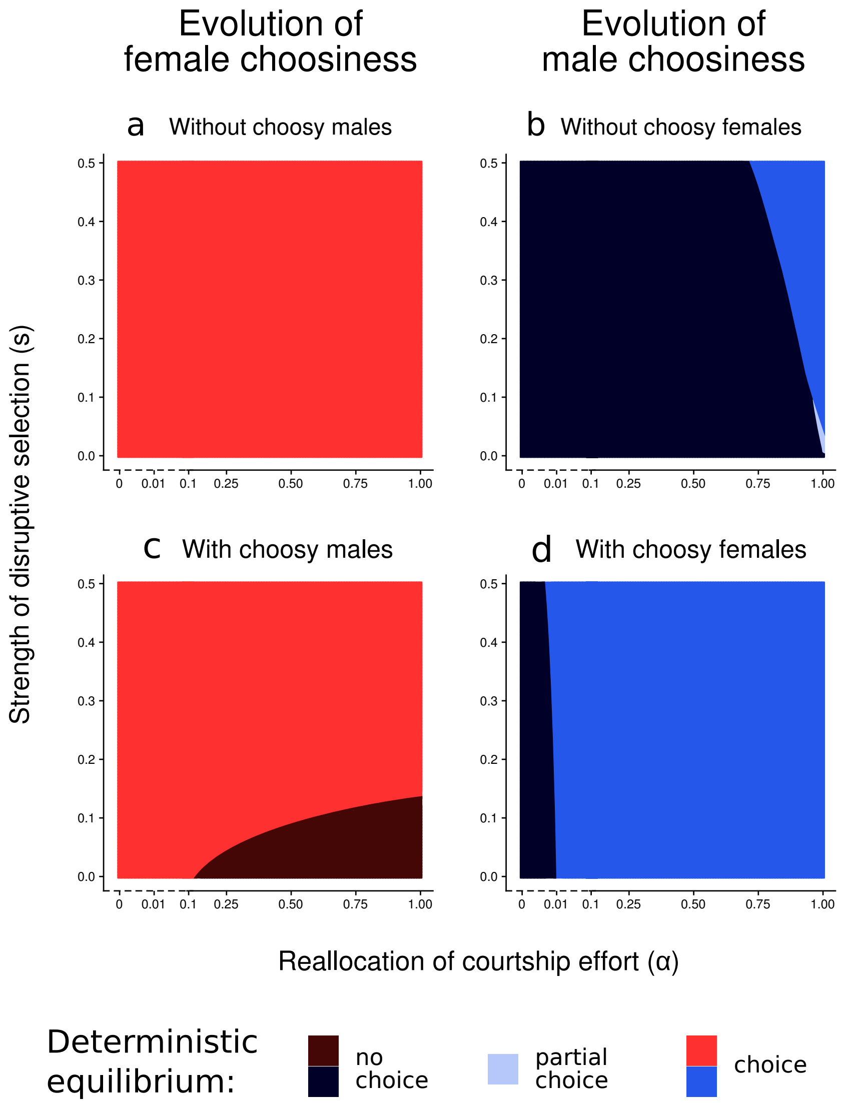


**Supplementary Figure 4:** Deterministic equilibrium **if choosiness can only evolve in one sex**. We vary the strength of disruptive selection ($s$), the extent of reallocation of courtship effort ($\alpha$) and the choosiness of the other sex. Without male choosiness, female choosiness is always favoured by indirect selection (a). Even without disruptive viability selection ($s = 0$), female choosiness is favoured by indirect sexual selection that acts on the ecological locus (a). Note that strong choosiness directly evolves in our model. A strong linkage disequilibrium develops between FF genotypes and AA or aa genotypes, such that choosy females are mostly homozygous at the ecological locus, increasing the mating success of homozygous males and indirectly favouring ecological divergence. Therefore, sexual selection does not inhibit ecological divergence in our model; this would not be the case if mating was initially random (such that intermediate phenotypes had the highest mating success) and if choosiness was evolving in small steps. If choosy males completely reallocate courtship effort towards preferred females, male choosiness is not necessarily favoured by indirect viability selection and may remain at an intermediate frequency (partial choice) (b). Indeed, male choosiness brings about competitive costs through distortion of male-male competition; choosy males often face stiffer competition than nonchoosy males. The evolution of male choice therefore involves some evolutionary pressures fundamentally different from those that determine the evolution of female choice. This has already been acknowledged by previous theory on male mate choice. In our model, we consider that reallocation of courtship effort towards preferred females can be partial. Not surprisingly, if choosy males do not reallocate courtship effort completely ($\alpha < 1$), strong viability selection is required for male choosiness to evolve (b). Choosiness in the other sex can change the deterministic outcome. First, female choosiness always favours the evolution of male choosiness (d). Second, male choosiness inhibits the evolution of female choosiness if viability selection is weak (c). This is caused by the selective forces represented in Fig. 1c.

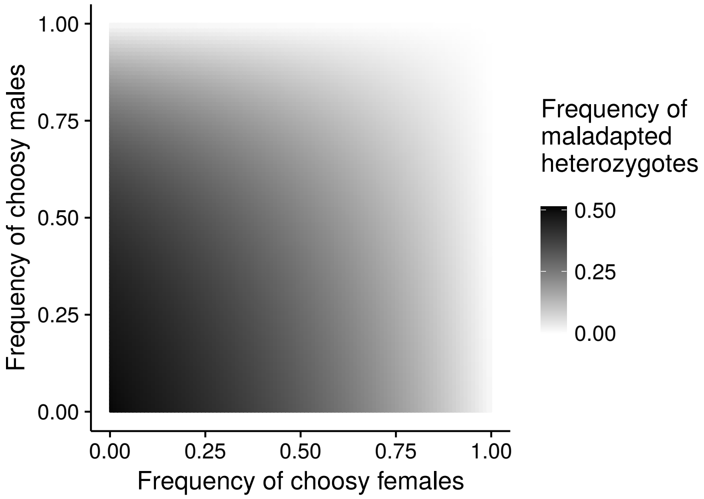


**Supplementary Figure 5:** Hybridization rate. We represent the deterministic frequencies of maladapted heterozygotes at the ecological locus (inversely proportional to the strength of reproductive isolation) in populations with given frequencies of choosy females and choosy males. Female and male choosiness synergistically reduce hybridization rate between ecotypes because we assume that choosy individuals do not completely avoid courting/mating across ecotype boundaries ($\epsilon_{\mathrm{m}} \neq 0$ and $\epsilon_{\mathrm{f}} \neq 0$). $s = 0.2$, $\alpha = 1$.

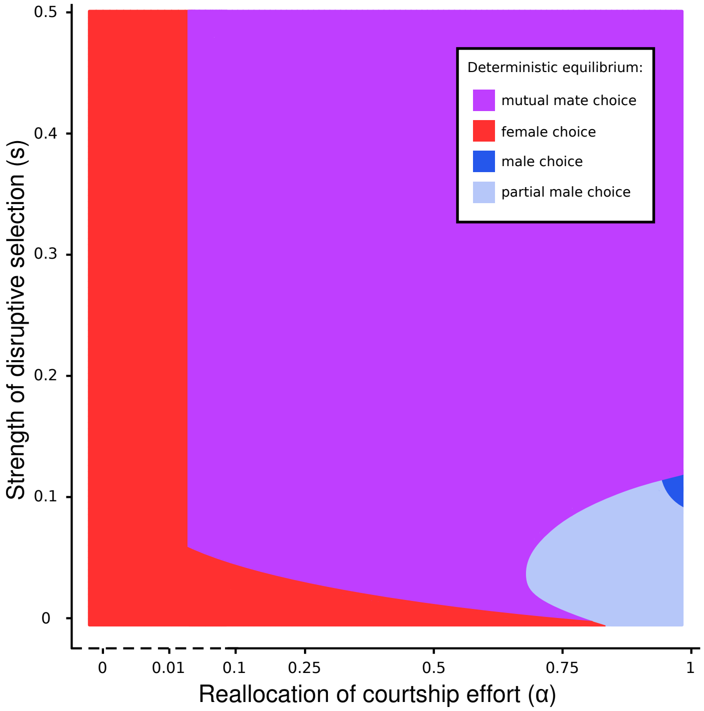


**Supplementary Figure 6:** Deterministic equilibrium for different combinations $(\alpha, s)$ of courtship reallocation and strength of disruptive viability selection **when alleles coding for choosiness are dominant**. See caption of Fig. 2 for details. Deterministic equilibrium is qualitatively similar to the one obtained with recessive choosiness alleles (Fig. 2a). Quantitative differences rely on the initial conditions. We implement initially 1% of choosy individuals. Therefore, initial frequencies of alleles F and M differ according to the dominance hierarchy implemented, and so does the deterministic equilibrium reached (in cases with more than one stable equilibrium).

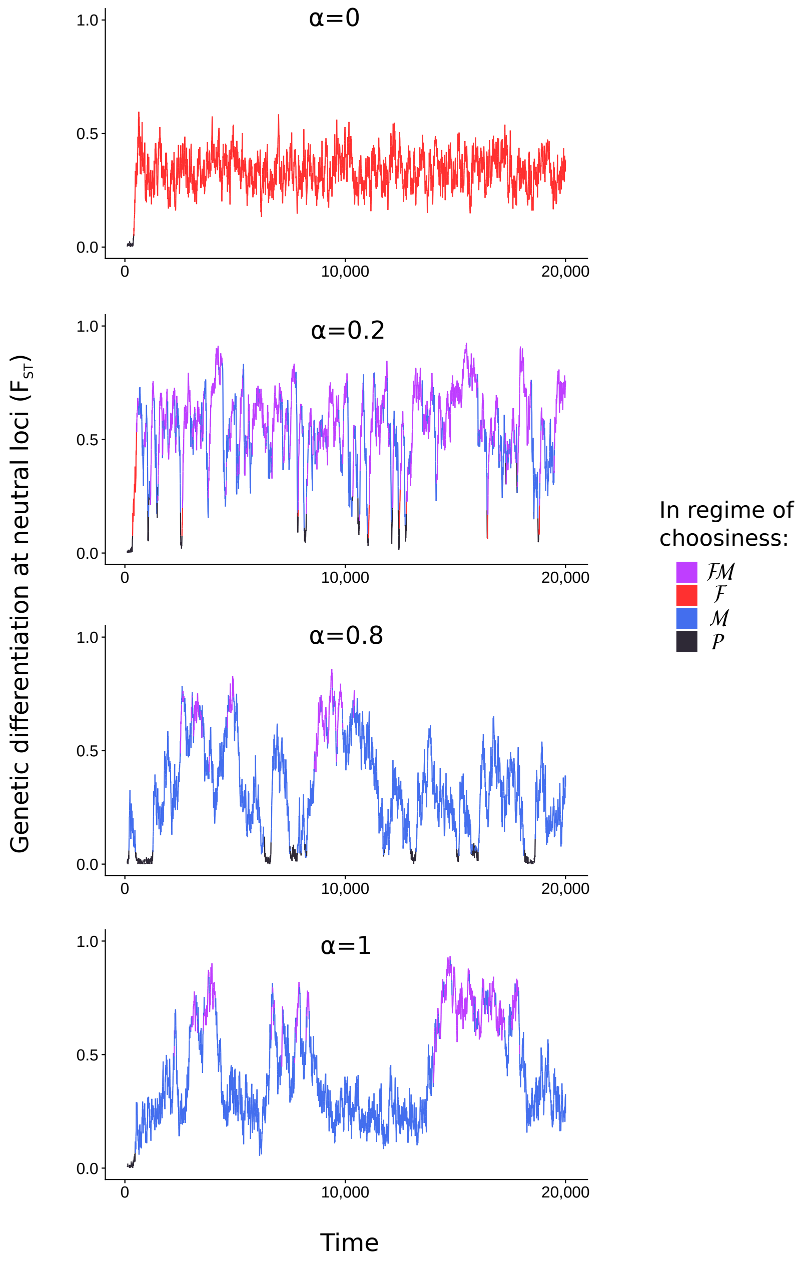


**Supplementary Figure 7:** Genetic differentiation at neutral loci. Here, we model the genetic differentiation of 20 additional unlinked diploid neutral loci (with 200 possible alleles per locus) among ecotypes AA an aa. In offspring, each neutral allele can mutate with probability $10^{-3}$ to any other allele. We compute the $F_{\mathrm{ST}}$ statistic as a measure of neutral genetic differentiation between ecotypes: $F_{\mathrm{ST}} = \frac{\Pi_s - \Pi_d}{\Pi_s}$. $\Pi_d$ and $\Pi_s$ represent the average number of pairwise differences in neutral loci between two individuals sampled from different ecotypes ($\Pi_d$, AA vs. aa) or from the same ecotype ($\Pi_s$, AA vs. AA or aa vs. aa). Simulations are run with different $\alpha$ values. In the graphs, the colour of the lines represent the regime of choosiness at each time step. Overall, male and female choosiness favour genetic differentiation among ecotypes at neutral loci ($F_{\mathrm{ST}} > 0$). If female mate choice evolves alone ($\alpha = 0$), genetic differentiation is stable at an intermediate level. $F_{\mathrm{ST}}$ does not reach 1 because female choice is subject to errors ($\epsilon_{\mathrm{f}} \neq 0$) and hybridization occurs at a small rate. If mutual mate choice is favoured ($\alpha > 0.01$), coevolutionary dynamics of female and male preferences temporarily homogenize populations ($F_{\mathrm{ST}}$ close to 0). Therefore, under selection for mutual mate choice, reproductive isolation is not stable.

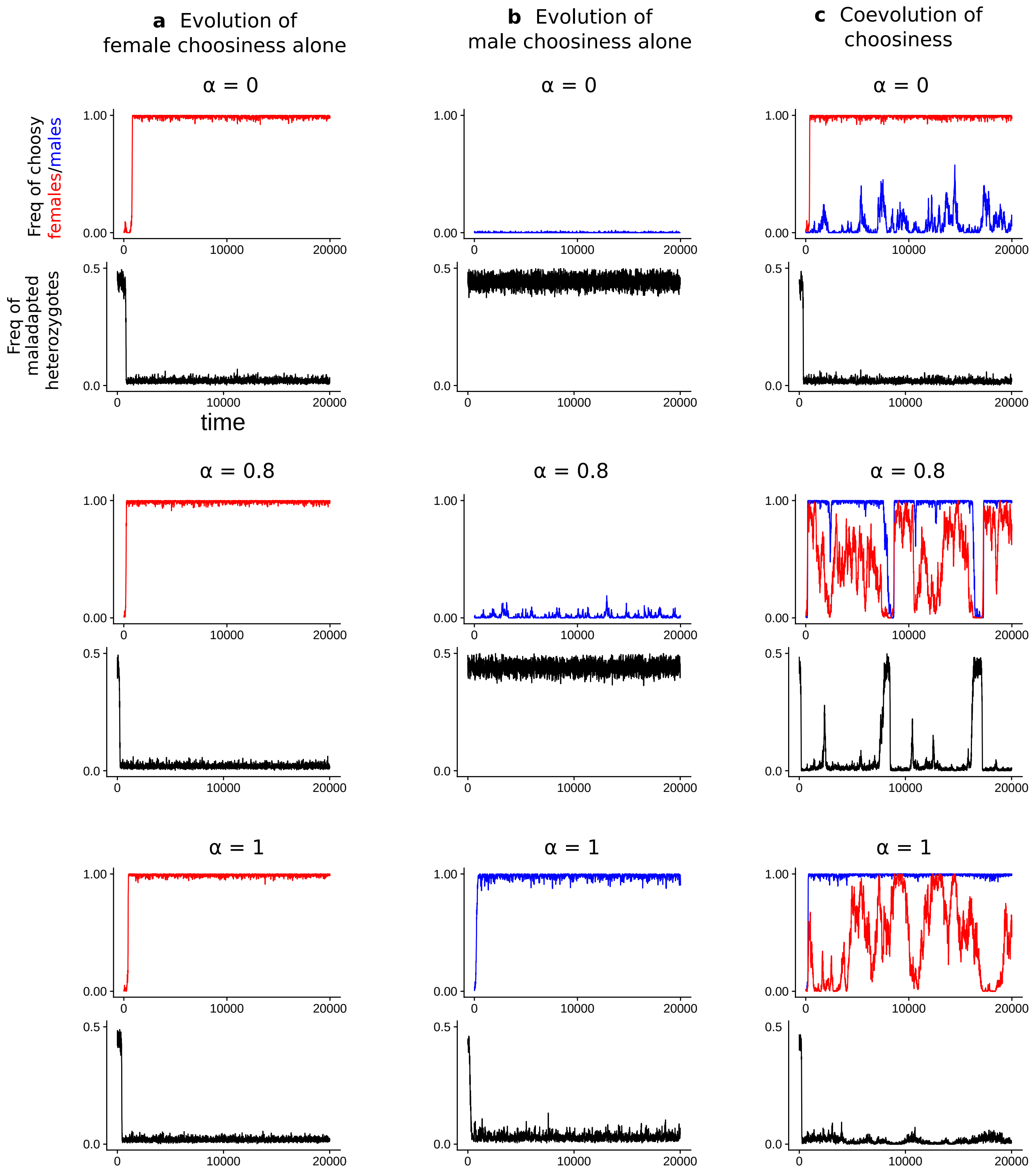


**Supplementary Figure 8:** Evolutionary dynamics of choosiness **if choosiness can only evolve in one sex** (a, b) **or in both sexes** (c) in stochastic simulations and resulting frequencies of maladapted heterozygotes at the ecological locus before viability selection ($s = 0.2$, $K = 500$). Frequencies of choosy females and choosy males are represented in red and blue, respectively. We vary the extent of reallocation of courtship effort ($\alpha$). If choosiness can evolve in one sex only, choosiness at equilibrium corresponds to choosiness obtained in deterministic simulations (Supplementary Fig. 4) and associates with stable reproductive isolation – i.e., reduced frequency of maladapted heterozygotes Aa is stable when choosiness evolves (a,b). If choosiness evolves in both sexes, the resulting coevolutionary dynamics can destabilize reproductive isolation (c).

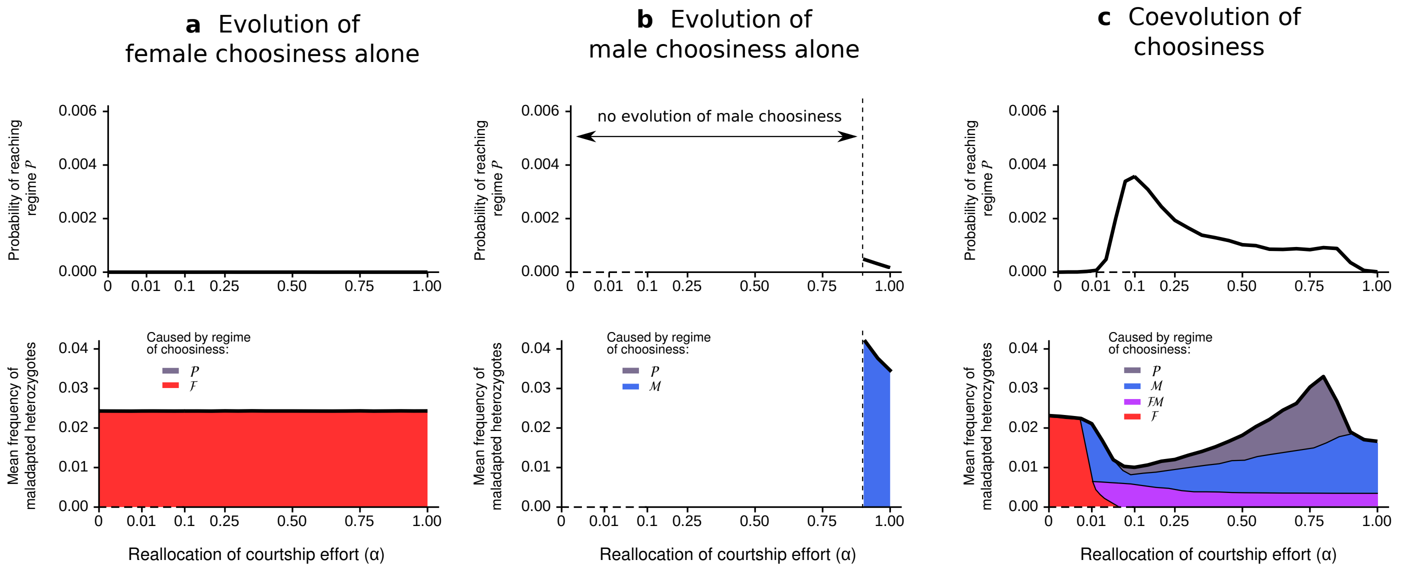


**Supplementary Figure 9:** Evolutionary dynamics of choosiness and resulting hybridization rate in stochastic simulations **if choosiness can only evolve in one sex** (a,b) **or in both sexes** (c) ($s = 0.2$, $K = 500$). We vary the extent of reallocation of courtship effort ($\alpha$). See caption of Fig. 5 for details. If choosiness can evolve in one sex only, choosiness at equilibrium corresponds to choosiness obtained in deterministic simulations (Supplementary Fig. 4) and associates with stable reproductive isolation – i.e., reduced frequency of maladapted heterozygotes Aa is stable when choosiness evolves (a,b). Note that we do not represent the mean hybridization rate = 0.5 when male choosiness does not evolve (for $\alpha \in [0, 0.9]$) (b). If choosiness evolves in both sexes, the resulting coevolutionary dynamics favours the evolution of male mate choice (even for low $\alpha$ values) but destabilizes reproductive isolation (c).

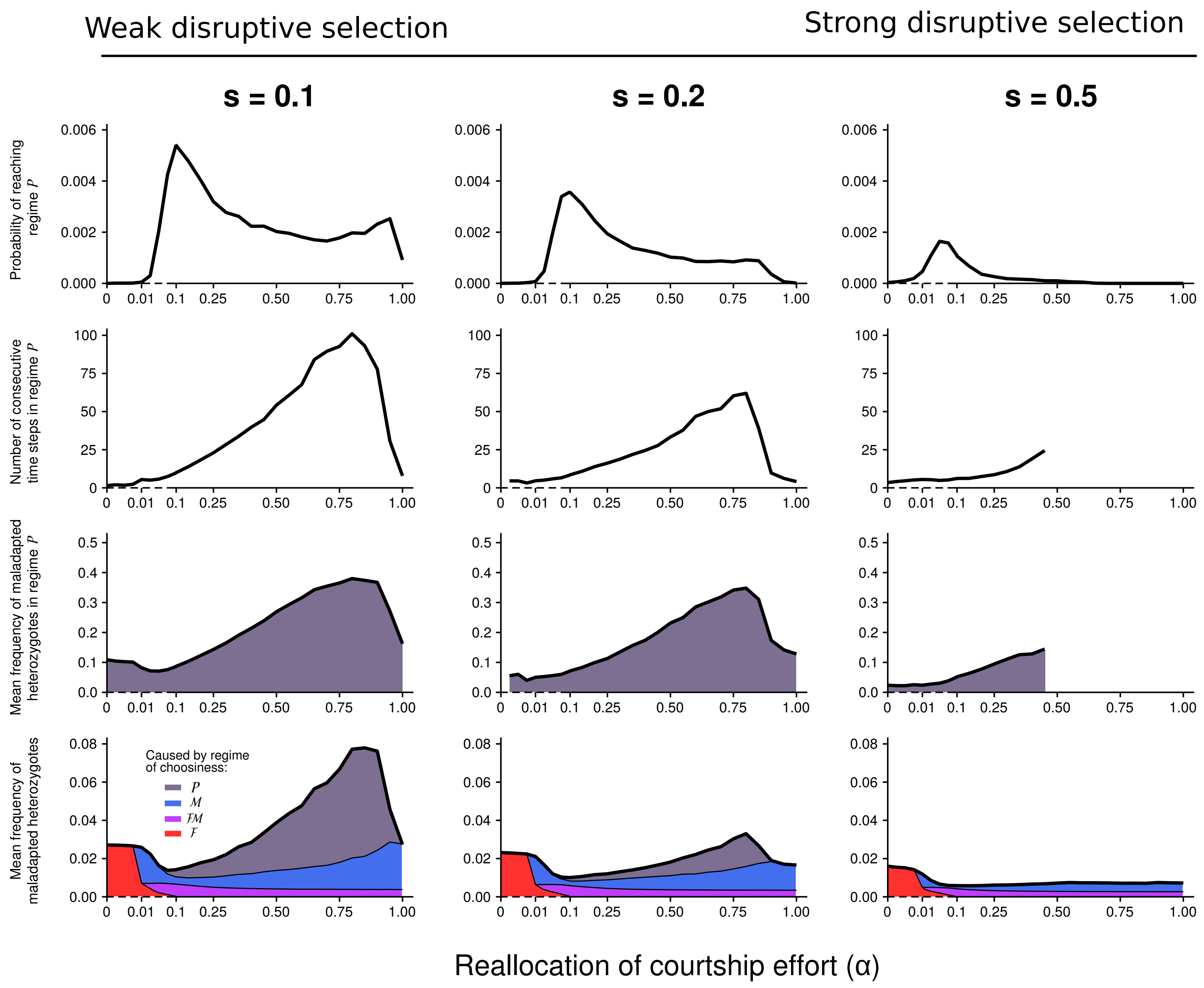


**Supplementary Figure 10:** Effect of the **strength of disruptive viability selection** on the coevolutionary dynamics of female and male choosiness and on the resulting hybridization rate in stochastic simulations ($K = 500$). We vary the strength of disruptive viability selection ($s = 0.1$, 0.2 or 0.5). See caption of Fig. 5 for more details. Female choosiness rarely evolves away from deterministic equilibrium in simulations with strong disruptive viability selection ($s = 0.5$) (small blue area). Likewise, preference cycling never occurs if disruptive viability selection is strong (no gray area). In that case, indirect viability selection favouring female and male choosiness is too strong for genetic drift to trigger preference cycling.

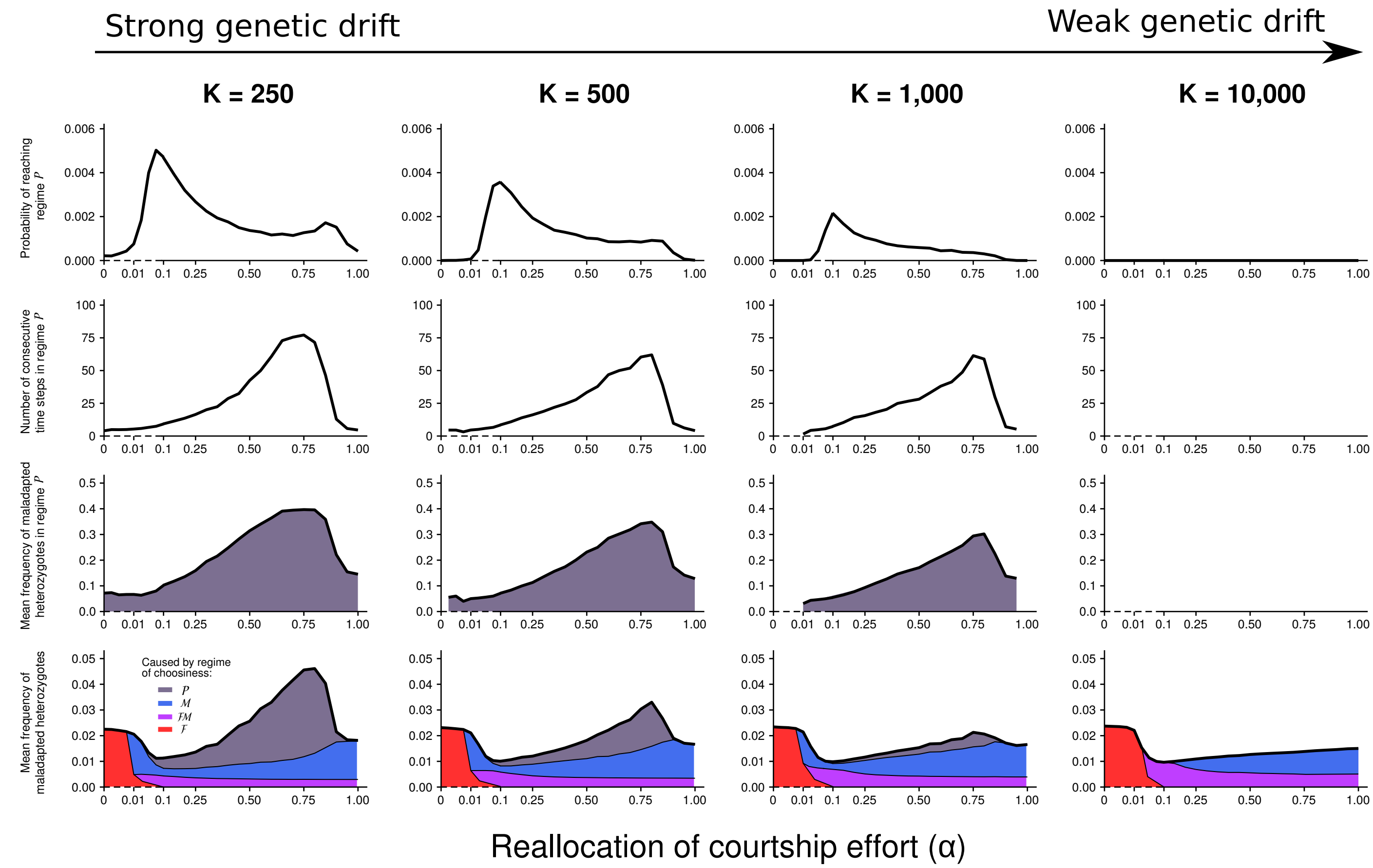


**Supplementary Figure 11:** Effect of the **carrying capacity** on the coevolutionary dynamics of female and male choosiness and on the resulting hybridization rate in stochastic simulations ($s = 0.2$ with $K = 250$, $500$, $1,000$ or $10,000$). See caption of Fig. 5 for more details. Here strong mutual mate choice is not maintained for $\alpha > 0.1$, even if drift is extremely weak ($K = 10,000$) (blue area in subfigures at the bottom). When males are choosy, selection favouring female choosiness is sufficiently weak for drift to occur. On the contrary, preference cycling does not occur if drift is weak (e.g., $K = 10,000$) (no gray area), as female choosiness rarely (almost never) drops to values low enough to change the direction of selection on male choosiness.

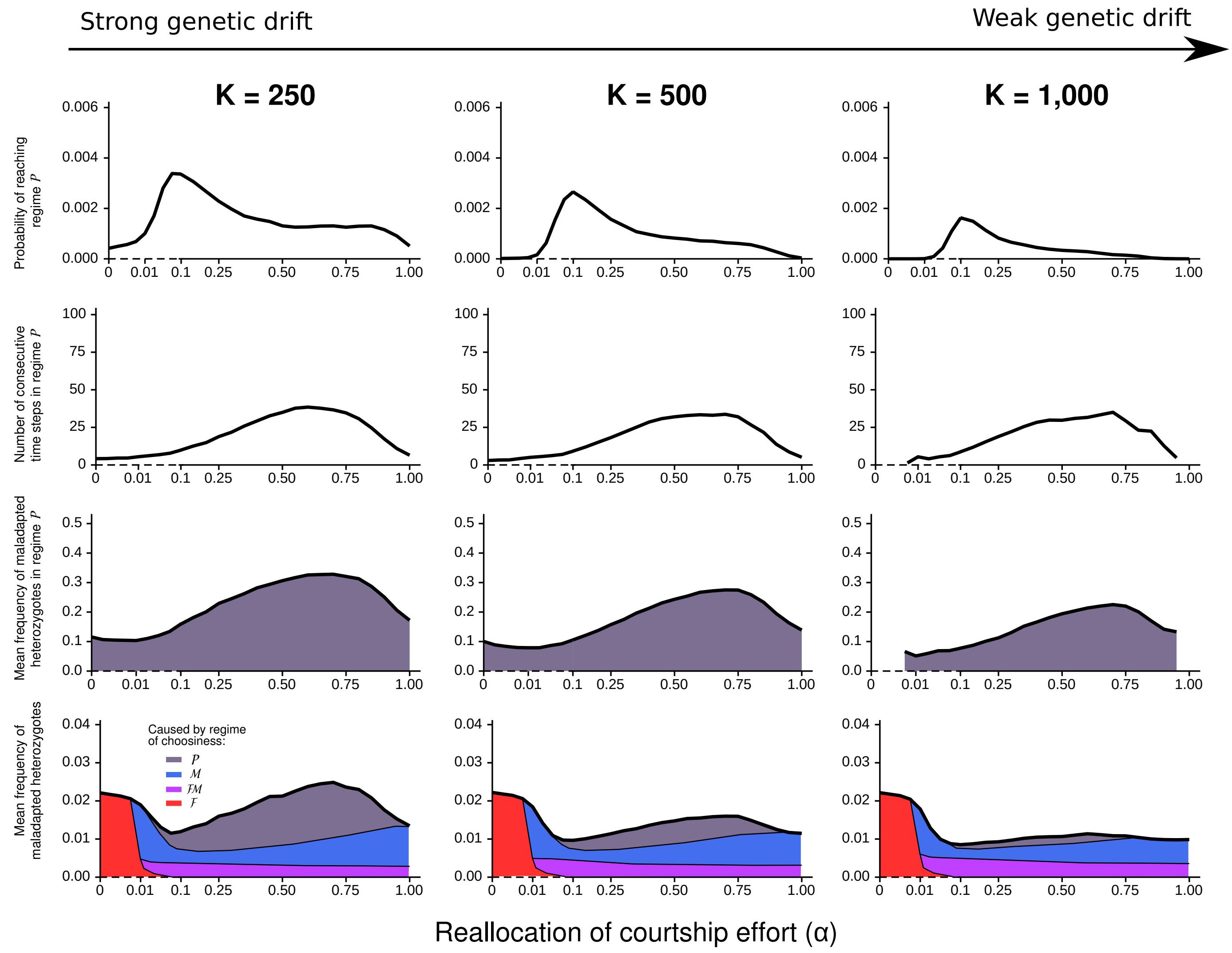


**Supplementary Figure 12:** Effect of the carrying capacity on the coevolutionary dynamics of female and male choosiness and on the resulting hybridization rate in stochastic simulations ($s = 0.1$ with $K = 250$, 500 or $10,000$) **when alleles coding for choosiness are dominant**. See caption of Fig. 5 for more details. Preference cycling can occur but is less likely than when alleles coding for choosiness are recessive (here, viability selection is weak; $s = 0.1$). With dominant choosiness alleles, once advantageous choosiness alleles have reached a high frequency, recessive nonchoosiness alleles are necessarily rare and thus mostly present in heterozygotes, such that complete fixation of the dominant choosiness allele is difficult. Given that the final approach to fixation of the male choosiness allele is less rapid, more nonchoosy males remain in the system. Therefore, in stochastic simulations, reproductive isolation is rarely nearly perfect if alleles coding for choosiness are dominant. This is not a condition favouring preference cycling. However, when preference cycling occurs, it strongly increases hybridization rate (with up to 30% of hybridization in regime $\mathcal{P}$; third row of subfigures).

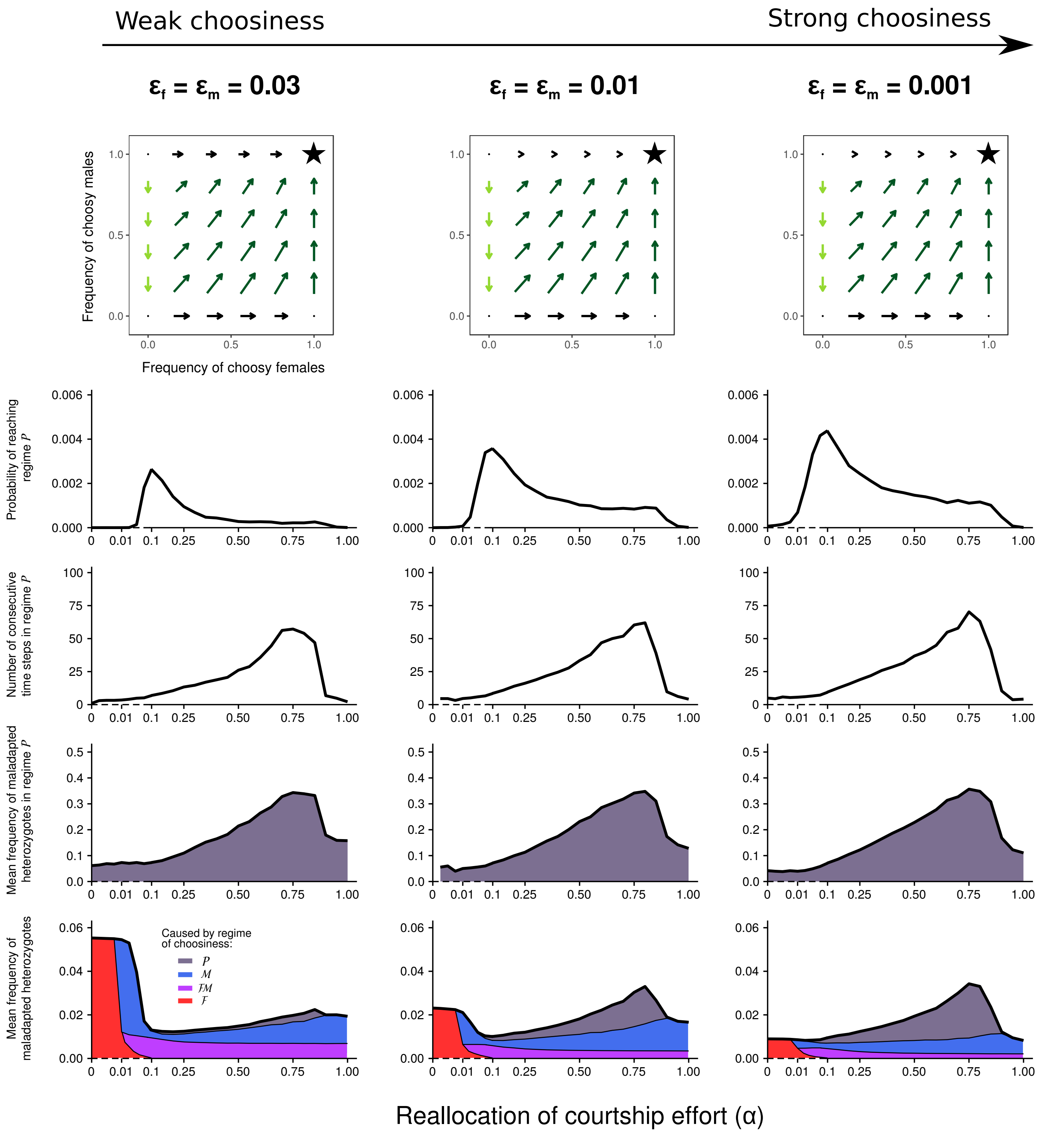


**Supplementary Figure 13:** Effect of the **strength of choosiness** on the coevolutionary dynamics of female and male choosiness and on the resulting hybridization rate in stochastic simulations ($s = 0.2$, $K = 500$). The top graphs are the selection gradients for $\alpha = 0.8$ (see caption of Fig. 4 for details). We vary the strength of choosiness (associated with genotypes MM and FF) ($\epsilon_f = \epsilon_m = 0.03$, $0.01$ or $0.001$). See caption of Fig. 5 for more details on the graphs at the bottom. For high $\alpha$, preference cycling rarely occurs if male choosiness is imperfect ($\epsilon_m = 0.03$) (little gray area) (see the cases with $\epsilon_f \neq \epsilon_m$ in Supplementary Fig. 14). In that case, female choosiness is strongly favoured by selection (cf selection gradients), and highly erroneously expressed male choosiness has the counterintuitive effect of decreasing the overall mean hybridization rate.

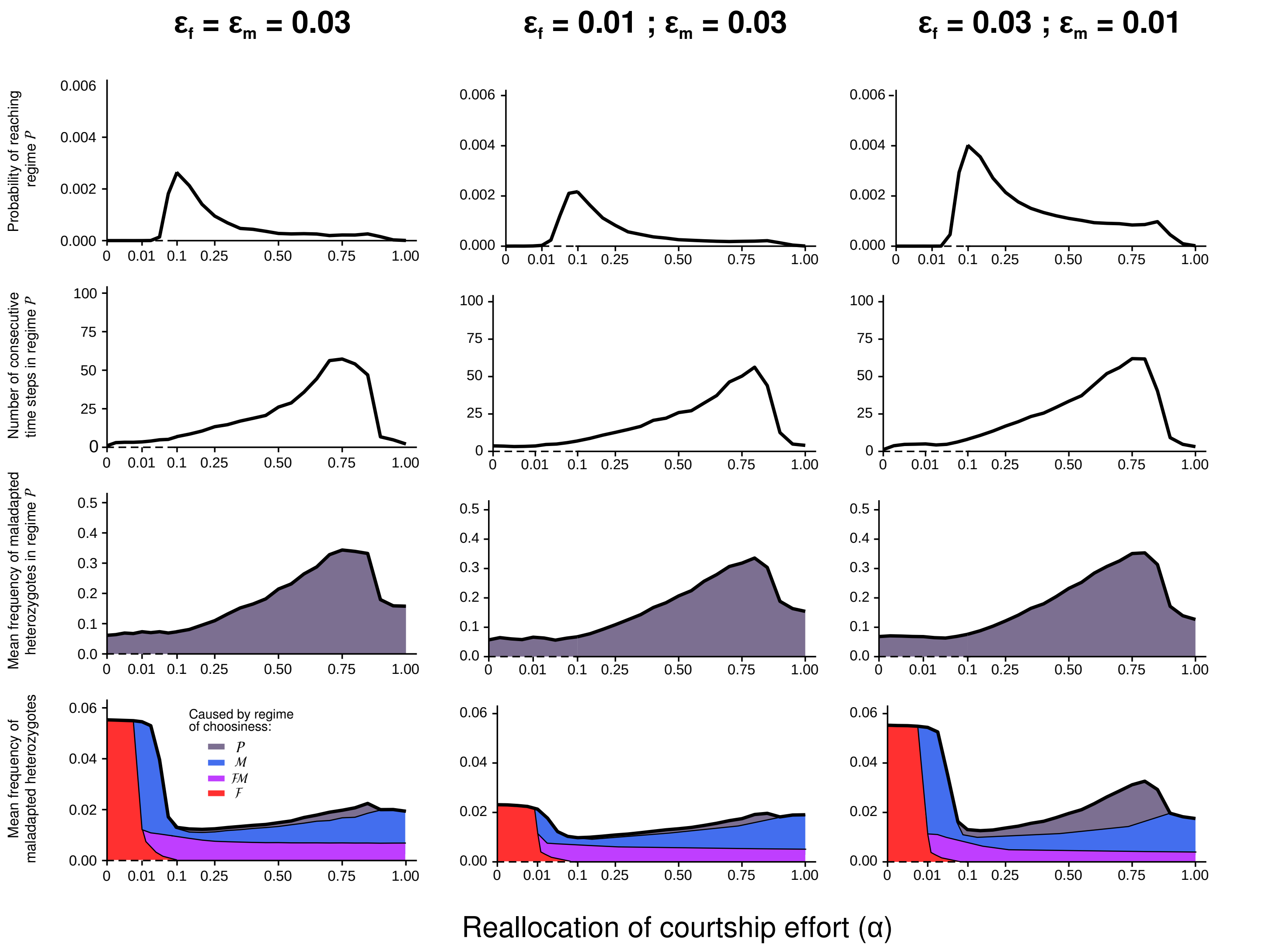


**Supplementary Figure 14:** Effect of the **sex-specific strength of choosiness** on the coevolutionary dynamics of female and male choosiness and on the resulting hybridization rate in stochastic simulations ($s = 0.2$, $K = 500$). We vary the strength of choosiness (associated with genotypes MM and FF) ($\epsilon_{\mathrm{f}}$, $\epsilon_{\mathrm{m}}$). See caption of Fig. 5 for more details on the graphs at the bottom. As shown in Supplementary Fig. 13, for high $\alpha$, preference cycling rarely occurs if male choosiness is imperfect (when $\epsilon_{\mathrm{m}} = 0.03$; little gray area). In that case, female choosiness is strongly favoured by selection, and preference cycling is rarely triggered by genetic drift. On the contrary, if only female choosiness is imperfect, preference cycling occurs and increases hybridization rate ($\epsilon_{\mathrm{f}} = 0.03$ ; $\epsilon_{\mathrm{m}} = 0.01$).

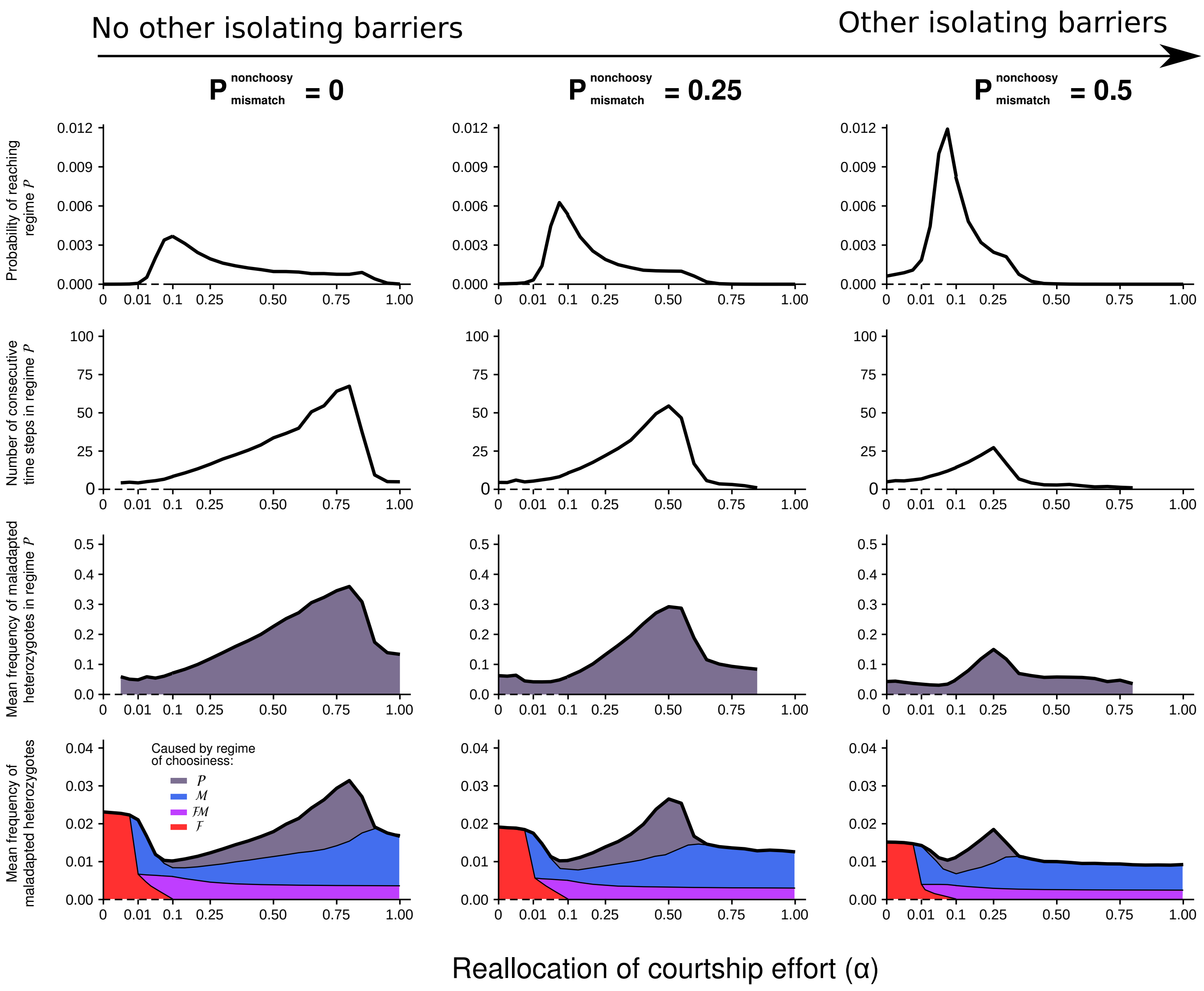


**Supplementary Figure 15:** Effect of **additional isolating barriers** on the coevolutionary dynamics of female and male choosiness and on the resulting hybridization rate in stochastic simulations ($s = 0.2$, $K = 500$). In these simulations, $P^{\male}_{m,f}$ and $P^{\female}_{f,m}$ of nonchoosy individuals are equal to a value $P^{\text{nonchoosy}}_{\text{mismatch}}$ if the genotypes m and f correspond to a mismatch at the ecological locus. In other words, we consider cases where assortative mating occurs even with only nonchoosy individuals in the system; the level of basal assortative mating is equivalent to the one driven by individuals expressing choosiness of strength $P^{\text{nonchoosy}}_{\text{mismatch}}$. See caption of Fig. 5 for more details. Preference cycling can occur if premating isolation leads to nearly complete reproductive isolation among ecotypes – e.g., if choosiness is nearly perfect (with up to 35% of hybridization in regime $\mathcal{P}$, left graph) or if other barriers already contribute to reproductive isolation (with up to 20% of hybridization in regime $\mathcal{P}$, right graph). By setting aside preference cycling, those conditions are the ones under which speciation appears the most likely.

### Supplementary Note 1 - Cost of female choosiness

In the main text, we assume that all females are guaranteed to mate. Yet, choosiness in females can also be costly. For instance, if population density is low, choosy females rejecting unpreferred males may remain unmated. In our model, we assumed that the mating success of females with and without a preference is equal. Here, by relaxing this assumption, we aim at testing if implementing costs of female choosiness can change the coevolutionary dynamics of female and male choosiness. Instead of the Equation 3 of the manuscript, we calculate the overall proportion of matings that occur between males of genotype $m$ and females of genotype $f$ as follows:

$$M_{m,f} = \widehat{C}_{m,f} \overbrace{\left( \overbrace{P^{♀}_{f,m}}^{\substack{\text{Baseline mating share}\\ \text{of a female of genotype } f \text{ with}\\ \text{a male of genotype } m}} + \beta \overbrace{\sum_{m'} \widehat{C}_{m',f}(1 - P^{♀}_{f,m'})}^{\substack{\text{Mating share that}\\ \text{a female of genotype } f\\ \text{reallocates}}} \times \overbrace{\frac{P^{♀}_{f,m}}{\sum_{m'} \widehat{C}_{m',f} P^{♀}_{f,m'}}}^{\substack{\text{Proportion of mating share that}\\ \text{a female of genotype } f \text{ reallocates}\\ \text{towards a male of genotype } m}} \right)}^{\text{Mating share of a female of genotype } f \text{ with a female of genotype } m} \quad (1)$$

with:

$$\widehat{C}_{m,f} = \frac{C_{m,f}}{\sum_{m'} C_{m',f}}.p_f \quad (2)$$

Parameter $\beta$ is analogous to $\alpha$ and reflects the extent of reallocation of mating share towards preferred males. If $\beta = 1$, all females are mating at the same rate and the mating success of females with and without a preference is equal ($\sum_{m'} \widehat{C}_{m',f} = p_f$). This corresponds to the idealized polygyny scenario we modelled in the main analyses. If $\beta = 0$, choosy females do not reallocate mating share towards preferred males. For instance, this corresponds to the scenario where females only encounter one potential mate in their lifetime, i.e., female choosiness is very costly. Since the elements of $M_{m,f}$ do not necessarily sum to one for $\beta < 1$, we normalize the genotype frequencies in offspring after recombination, such that $\sum_i p_i = 1$.

If costs of female choosiness are high (many lost mating opportunities, $\beta < 0.85$), female choosiness does not evolve even under strong disruptive viability selection ($s = 0.2$) and in the absence of male choosiness. Consequently, male choosiness does not evolve either (because in the absence of female choosiness, it is disfavoured by direct sexual selection) and the population remains in a state of random mating (Supplementary Fig. 16c-d).

If costs of female choosiness are weak ($\beta > 0.85$), coevolutionary dynamics of female and male choosiness are qualitatively similar to those without costs of choosiness (Supplementary Fig. 16b-c vs. Fig. 4a-d). In particular, even if female choosiness is costly, preference cycling is still induced by genetic drift and completed by selection.

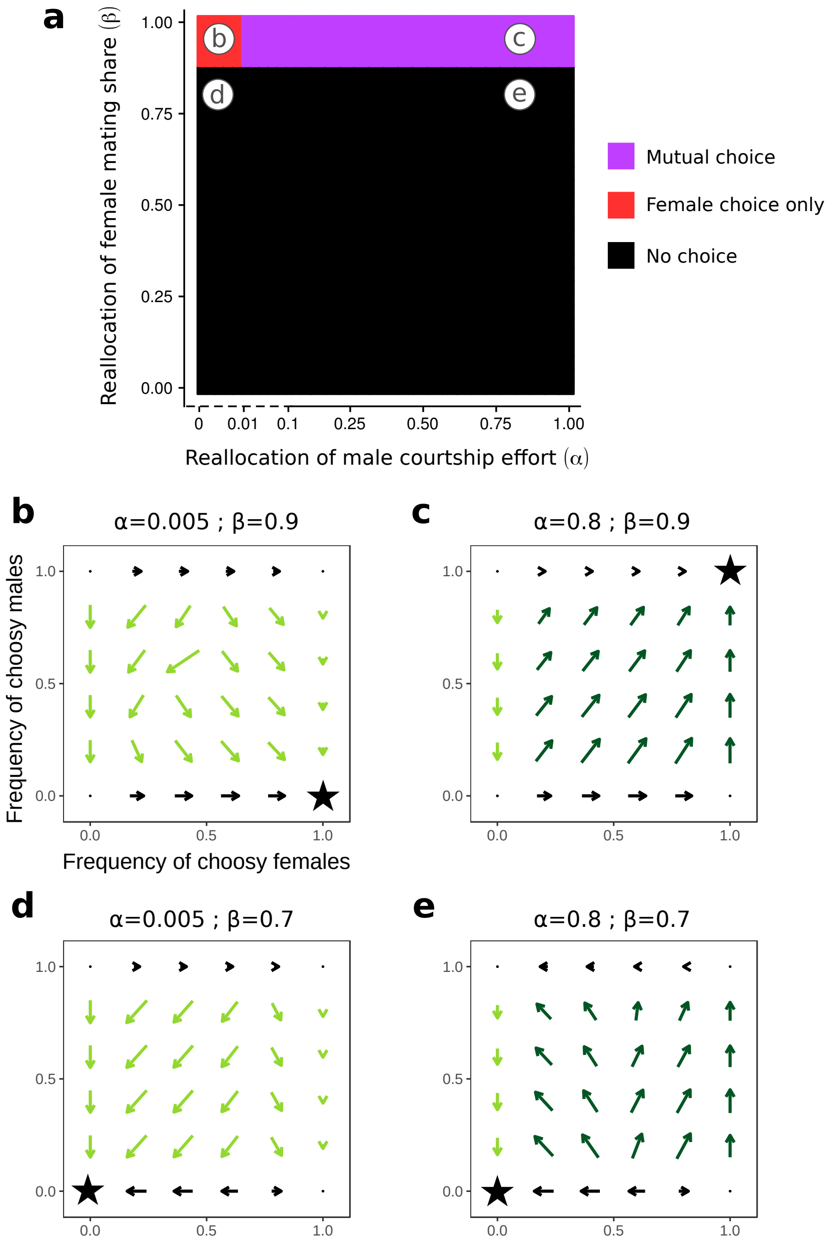


**Supplementary Figure 16:** Deterministic equilibrium for different combinations $(\alpha, \beta)$ of male courtship reallocation and female mating share reallocation (a). Frequencies of choosy males and females either reach one or zero, and we describe the regime of choosiness at equilibrium accordingly. Subfigures b-e are deterministic selection gradients for female and male choosiness and refer to specific combinations of $(\alpha, \beta)$ marked in the subfigure a. See caption of Fig. 4 for details. In particular, preference cycling is not more prevalent if female choosiness is weakly costly (b,c) ; the initial decline of female choosiness still needs to be induced by genetic drift, not by selection against costly choice (cf. the direction of the arrows in the selection gradients). $s = 0.2$.

### Supplementary Note 2 - Continuous choosiness

For simplicity, we only included two different alleles per locus in our population genetic model, and choosiness is therefore a binary trait (nonchoosy vs. choosy genotypes). Yet, intermediate levels of choosiness are observed in nature. To account for continuous choosiness, we here place our model in the adaptive dynamics framework.

We assume female and male choosiness are discretized into 10 distinct levels, each of which is coded by a distinct allele. Namely, we implement alleles $F_1$, $F_2$,..., $F_{10}$ at the $F$ locus. $F_1$ is coding for nonchoosiness ($\epsilon_f = 1$) and $F_{10}$ for complete choosiness ($\epsilon_f = 0.01$). $F_2$, $F_3$,... $F_9$ are coding for intermediate choosiness ($\epsilon_f = 0.9$, 0.8, ... 0.1, respectively). Like in our diploid population genetic model, we assume that alleles coding for a choosier state are recessive. The same applies at locus $M$.

Following the adaptive dynamics approach, we assume that choosiness loci can only mutate to the closest neighboring alleles (stepwise-mutation model). For instance, a population at regime $F_2F_2$ can only be invaded by mutants carrying allele $F_1$ or allele $F_3$. We first determine numerically the probability of invasion for each allele, using our diploid population genetic model (stochastic simulations). The invasion of an allele coding for a choosier or less choosy state depends on female and male choosiness in the population (Supplementary Fig. 17) and on the carrying capacity $K$ (not shown).

Using these invasion probabilities, we simulate the successive invasions of mutant alleles assuming that only one mutation can occur at a time (Supplementary Fig. 18). Female mate choice and mutual mate choice evolve for $\alpha = 0$ and $\alpha = 1$, respectively. 'Preference cycling' occurs for intermediate $\alpha$ (= 0.5), just like in our model without continuous choosiness. When female choosiness drifts below a threshold, selection favouring indiscriminate male courtship leads to a transient regime of random mating. Overall, these evolutionary dynamics with continuous preference traits are consistent with the stochastic simulations of our simple diploid population genetic model.

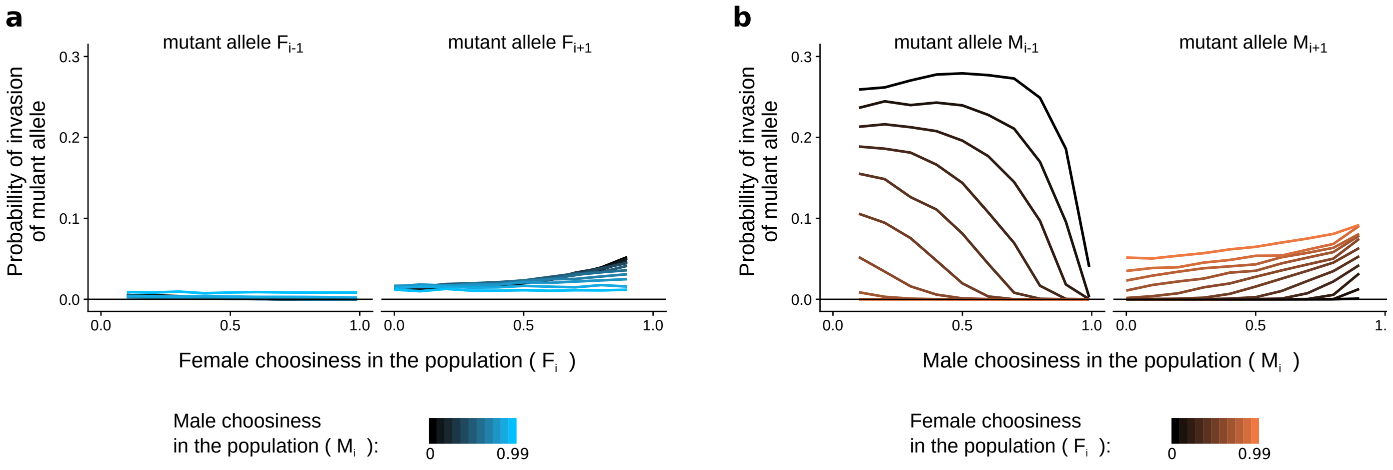


**Supplementary Figure 17:** Probability of invasion of mutant allele at locus $F$ (a) or at locus $M$ (b) estimated from stochastic simulations of our diploid population genetic model for $K = 500$, $s = 0.2$ and $\alpha = 0.5$. Neutral or deleterious alleles can get fixed by chance (e.g., fixation alleles $F_{i-1}$ when males are choosy). Probabilities of invasion of mutant alleles (i.e., of reaching a frequency $> 0.99$) depend on female and male choosiness in the population.

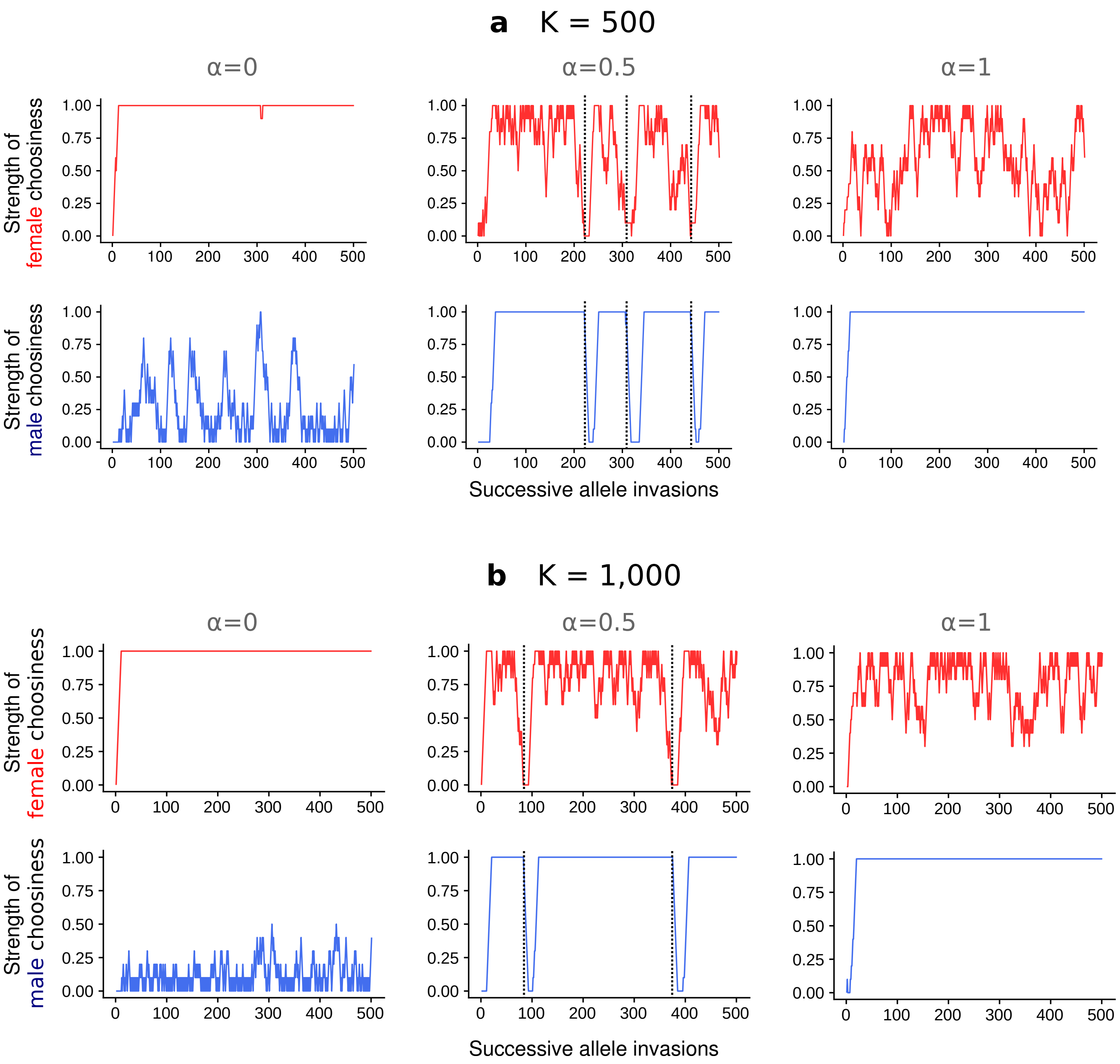


**Supplementary Figure 18:** Evolutionary dynamics of continuous female and male choosiness for different values of $K$ and $\alpha$. For $\alpha = 0.5$, when female choosiness drifts below a threshold, selection leads to a transient regime of random mating (dashed vertical lines). For $\alpha = 0$, partial male choosiness can evolve by chance because male choosiness is only weakly deleterious when females are choosy (cf. the selection gradient in Fig. 4a); however, selection is still strong enough to inhibit the evolution of strong male choosiness. $s = 0.2$.
